# Systematic identification of engineered methionines and oxaziridines for efficient, stable, and site-specific antibody bioconjugation

**DOI:** 10.1101/748160

**Authors:** Susanna K. Elledge, Hai L. Tran, Alec H. Christian, Veronica Steri, Byron Hann, F. Dean Toste, Christopher J. Chang, James A. Wells

**Author notes:** To whom correspondence should be addressed: James A. Wells, University of California, San Francisco, 1700 4^th^ Street, 504 Byers Hall MC2552 San Francisco, CA 94143.

## Abstract

Chemical modification of antibodies is one of the most important bioconjugations utilized by biologists and biotechnology. To date, the field has been dominated by random modification of lysines or more site-specific labeling of cysteines, each with attendant challenges. Recently we have developed oxaziridine chemistry for highly selective and efficient sulfimide modification of methionine called redox-activated chemical tagging (ReACT). Here, we systematically scanned methionines throughout one of the most popular antibody scaffolds, trastuzumab, for antibody engineering and drug conjugation. We tested the expression, reactivities, and stabilities of 123 single engineered methionines distributed over the surface of the antibody when reacted with oxaziridine. We found uniformly high expression for these mutants and generally good reaction efficiencies with the panel of oxaziridines. Remarkably, the stability to hydrolysis of the sulfimide varied more than ten-fold depending on temperature and the site of the engineered methionine. Interestingly, the most stable and reactive sites were those that were partially buried, likely because of their reduced access to water. There was also a ten-fold variation in stability depending on the nature of the oxaziridine, which we determined was inversely correlated with the electrophilic nature of the sulfimide. Importantly, the stabilities of the best analogs and antibody drug conjugate potencies were comparable to those reported for cysteine-maleimide modifications of trastuzumab. We also found our antibody drug conjugates to be potent in a breast cancer mouse xenograft model. These studies provide a roadmap for broad application of ReACT for efficient, stable, and site-specific antibody and protein bioconjugation.

## Introduction

Monoclonal antibodies are among the most universal tools in biology and medicine^1^. Chemical bioconjugation has been instrumental in expanding the utility of monoclonal antibodies, both as probes and therapeutics, by facilitating covalent attachment of a variety of moieties such as fluorophores^2^, metal chelators^3^, nucleic acids^4–6^, as well as toxins in the form of antibody drug conjugates (ADCs)^7–10^. ADCs have revolutionized the ability to selectively deliver cytotoxic compounds to cancer cells by binding to tumor-specific antigens. As a result, ADCs can be an improvement over standard chemotherapy treatment by simultaneously increasing targeting efficiency and reducing off-target toxicity^11–14^. There are currently five FDA approved ADCs and more than 100 clinical trials to develop new ADC therapies^11,12^.

The bioconjugation method is a critical consideration for any protein modification application. Ideally the modification should be efficient, stable, site-selective for homogeneity/reproducibility, and should not scar the overall functional properties of the protein^12^. To date, researchers have typically targeted lysine or cysteine residues for chemical conjugation due to the robustness and commercial accessibility of functionalized N-hydroxy succinamides to form stable amide bonds with lysines or maleimides to form stable thioether linkages to cysteines^11–14^. Three out of the five FDA approved ADCs target lysines for conjugation^14^. However, antibodies typically have about 40 surface exposed lysine residues per IgG which can result in more than one million different ADC species^12^. These conjugates are therefore highly heterogenous in terms of conjugation site and drug-to-antibody-ratio (DAR) that form a gaussian distribution usually ranging from zero to eight ^12^. The conjugated cytotoxic drugs tend to be hydrophobic causing aggregation, immunogenicity, faster clearance rates and thus differences in the pharmacodynamic properties of the conjugate^11,15^. Additionally, the specific site of conjugation can have an effect on the efficacy of the ADC based on the stability and the aggregation propensity of the resulting derivative^12^.

Cysteine is becoming more commonly used as it is far less abundant than lysine and affords greater site-selectivity. This approach usually involves reducing the interchain disulfide bonds of the antibody and re-conjugating to a thiol reactive moiety, either resulting in disrupted disulfide bonds or re-bridged disulfides^16^. However, disulfide reduction can still lead to heterogenous mixtures unless the reduction is site-specific^11^. Additionally, the reduction process can cause disulfide scrambling which can disrupt the stability and even the structure of the antibody by forming different disulfide connections^17^. More recently researchers have systematically explored the introduction of single cysteine residues into the therapeutic antibody, trastuzumab, to identify sites for stable and specific conjugation without affecting its binding functions^18,19^. Interestingly, the conjugate stability is highly dependent on the cysteine site for both disulfide and maleimide based conjugations^19^. Although lysine and cysteine modifications dominate the field, other conjugation strategies are being developed and utilized including enzymatic conjugation, glycan modification, and un-natural amino acid incorporation^11,14^. These strategies result in homogenous conjugates but can be limited in terms of DAR and involve introducing larger sequence scars, either by peptide motifs or altering natural glycosylation^14^. While there has been progress in site-specific conjugation technologies for ADCs and proteins in general, we believe there is a need to add and improve the armamentarium of chemical conjugation strategies for more efficient, stable, and site-selective bioconjugations.

Recently, a methionine specific chemistry has been developed, redox-activated chemical tagging (ReACT), to efficiently and site-specifically conjugate to methionine residues on proteins (**Fig. 1A**)^20^. The ReACT methionine chemistry involves oxidation of methionine to form a sulfimide adduct with an oxaziridine molecule functionalized with an alkyl-azide to allow cargo attachment via click chemistry. Methionine is the second least-abundant residue in proteins after tryptophan^21^, making it an ideal target to site-specific conjugation. Most methionine residues are buried and therefore inaccessible, making it a potentially excellent target for bioengineered chemical conjugation.

**Figure 1.**
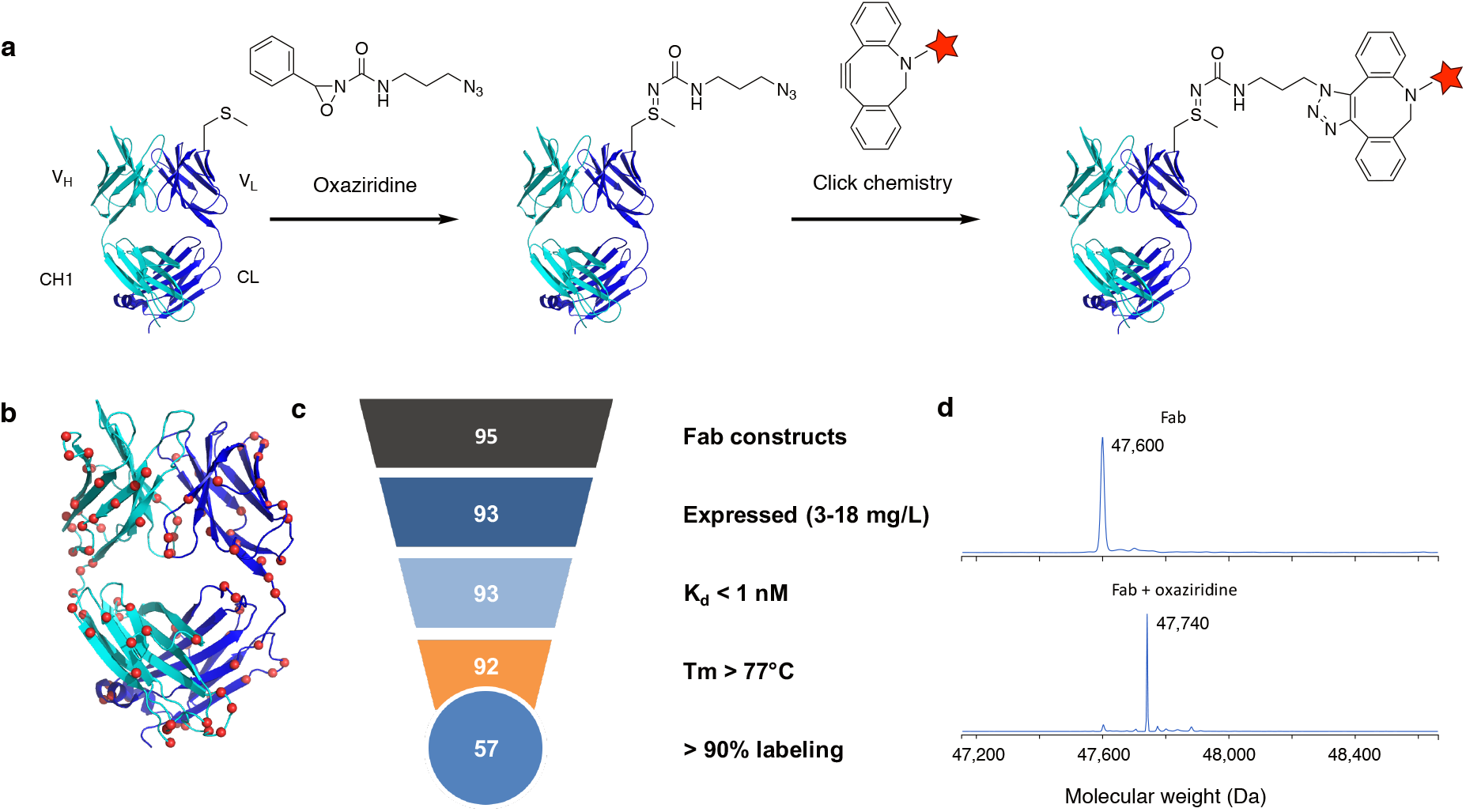
Oxaziridine labeling of most accessible sites on model αGFP-Fab in the trastuzumab scaffold. (a) Scheme of modular oxaziridine labeling on trastuzumab Fab. The Fab light chain is shown in dark blue and the heavy chain is shown in cyan. After conjugation with oxaziridine, different functionalities can be clicked on with a DBCO reagent. (b) The top 95 calculated accessible sites on the Fab scaffold are shown as red spheres. (c) Triage of the 95 most accessible mutants is shown. Each site was engineered to methionine on a model αGFP-Fab in the trastuzumab scaffold. Sites were then assessed for expression, affinity, structural stability, and labeling percentage. (d) Representative ESI mass spectra of labeling Fab with oxaziridine, shown by a mass shift of 140 (expected: 140).

To enable robust and expanded application of ReACT for bioconjugation, we methodically analyzed how the character of the engineered methionine site and oxaziradine analog affects reaction efficiency and stability of the antibody conjugate. We systematically scanned single methionine residues at 123 exposed or partially buried sites in the trastuzumab scaffold to identify sites for optimal conjugation. We found the engineered sites to be highly reactive and high yielding, resulting in site-selective conjugates without affecting antibody binding affinity. Surprisingly, we found large differences in stability to hydrolysis of the linkage based on the nature of the oxaziridine and the labeled site. We identified systematic factors that affect stability that we believe are portable to other antibodies and proteins. We found sites of equal stabilities to top thiol sites, and designed antibody drug conjugates with DAR of 2 with comparable potency in *in vitro* and *in vivo* models. These studies show that bioconjugation through methionine using ReACT is an outstanding approach for site-specific, high yielding and stable antibody derivative and identify critical factors for employing ReACT to proteins in general.

## Results

### High-throughput scan of the top 95 most accessible sites on the trastuzumab scaffold

We chose trastuzumab as the antibody scaffold of choice for our studies for a number of reasons. The trastuzumab framework is popular for humanization due to its high stability, high expression in mammalian cells, high developability, and broad use that is now utilized in parts of three different approved antibody drugs (trastuzumab, bevacizumab, and omalizumab) and the TDM-1 anti-Her2 ADC (ado-trastuzumab emantisine). Synthetic complementarity-determining region (CDR) libraries have been constructed on the trastuzumab scaffold^22^ and used by the Recombinant Antibody Network for industrialized recombinant antibody generation to over 500 protein targets^23^. The Fab arms in trastuzumab contain three methionines that are buried (**Supplementary Fig. 1)**. Indeed previous studies from our group showed these buried methionines to be unreactive to ReACT but when we attached a single Met to the C-terminus of the light chain we found it could be labeled quantitatively with a simple oxaziridine reagent and conjugated with a fluorophore^20^. While this site can be labelled quantitatively and could be useful for short-term *in vitro* studies, we found it becomes extensively (>80%) hydrolyzed over three days at 37°C **(Supplementary Fig. 2)** and thus is not suitable for long-term studies or ADC development.

To expand the use of ReACT for antibody bioconjugations we sought to systematically determine how methionine mutation, site of labeling, and compound nature affects expression, labeling efficiency, binding affinity, and stability of the antibody (**Fig 1A**). We first focused on exposed sites on a well characterized αGFP antibody built on the trastuzumab scaffold as a model for ease of assay^23^. We calculated the surface accessibility of the methionine sulfur for all possible surface methionine substitutions. We mutated the top 95 most accessible sites to methionine (**Fig. 1B**, **Supplementary Table 1**) and expressed each individual mutant in the αGFP Fab without mutating the three intrinsic and unreactive buried methionines. Remarkably, of those 95 sites, 93 methionine mutants expressed with high yield in *E. coli* (3-18mg/L). All 93 retained high binding affinity for GFP, and 92 of those retained high thermostability as measured by differential scanning fluorimetry (DSF). When tested for labeling with 5 equivalents of the oxaziridine reagent (oxaziridine **1**) for 2 hours, 57 mutants labeled to greater than 90% (**Fig. 1C**). This could potentially be improved with higher equivalents of oxaziridine. All mutants labeled stoichiometrically and specifically at the mutated methionine residue, as determined by whole protein mass spectrometry (**Fig. 1D**; **Table 1**). These data suggest tremendous flexibility in generating site specific methionine conjugations.

While these sites are likely useful for short-term studies such as immunofluorescence or other *in vitro* studies we wanted to test their suitability for longer term *in vivo* applications. Of the 57 highly labeled sites, we chose 12 representative sites to test conjugation stability as a function of location and temperature (**Fig. 2A**). The 12 candidate sites spanned both the heavy and light chain, as well as the variable and constant domains of the Fab arm. We incubated each methionine-oxaziridine conjugate at 4°C, 25°C, and 37°C for 3 days and measured the remaining conjugate by whole protein MS (**Fig. 2B**). We found a strong temperature dependence for hydrolysis from 4°C, 25°C, and 37°C. There was considerable variation among the sites, but all sites had less than 60% remaining conjugate after 3 days at 37°C. The product had a +16 mass shift consistent with hydrolysis of the sulfimide to a sulfoxide product, which has also been previously reported^24^. Since ADCs can have circulation times up to weeks in the body, it is essential that the linkage is stable for an extended period of time at biological temperatures to retain ADC potency and to eliminate off-target toxicity due to free drug release. Although these stabilities are sufficient for the many *in vitro* uses for antibody conjugation, we sought to extend the stability of the antibody conjugate for ADCs.

**Figure 2.**
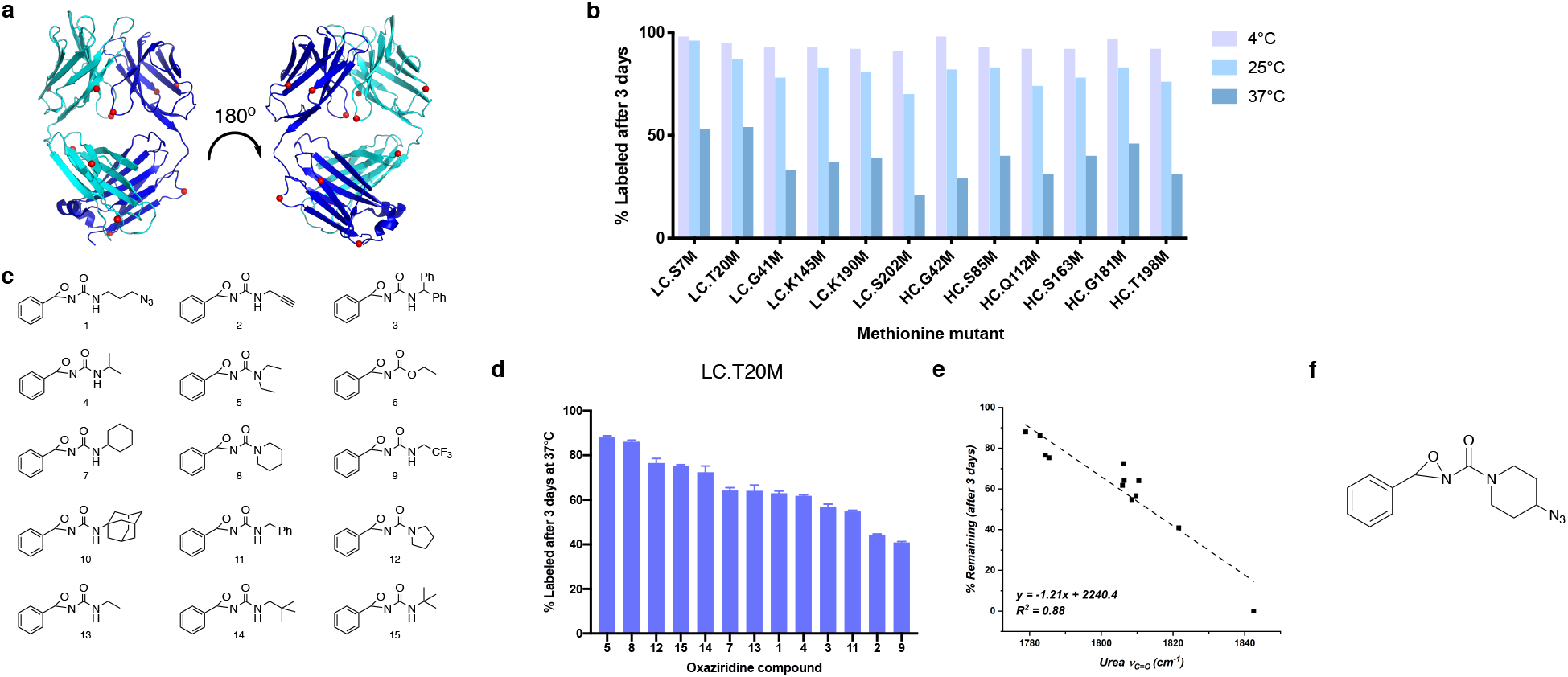
Labeling and stability of top 12 accessible sites with different oxaziridine compounds. (a) The trastuzumab Fab structure showing the top 12 labeled sites (>95%) as red spheres. (b) Conjugate stability of the top 12 sites labeled with oxaziridine at 4°C, 25°C, and 37°C over 3 days varies across sites and temperatures. These sites show a significant decrease in stability to hydrolysis at elevated temperatures. (c) Panel of oxaziridine derivatives tested for stability and (d) conjugate stability at site LC.T20M over 3 days at 37°C is depicted for each derivative (n=3). Oxaziridine **6** is not shown because it showed 0% stability. Oxaziridine **10** is not shown because no initial labeling could be detected. (e) Correlation of compound conjugate stability of LC.T20M and carbonyl stretching frequency (*ν*_C=O_). As the substitution on the oxaziridine/corresponding sulfimide becomes more electron rich, the less electrophilic the sulfimide becomes, thus increasing conjugate stability on the protein. (f) The structure of the piperdine-derived oxaziridine azide **8**.

### Enhancing stability of oxaziridine conjugates

We took two approaches to improve conjugation stability: (1) test different substituted oxaziridine analogs to improve linkage stability and (2) test more buried sites on the Fab scaffold that we hypothesized could better shield the sulfimide from hydrolysis. We obtained 15 different oxaziridine molecules with various functionalities appended to the urea group to determine if the resulting sulfimide bond could be further stabilized (**Fig. 2C**). We chose one representative site, LC.T20M, that showed moderate stability at 37°C for oxaziridine **1**. All compounds were conjugated to LC.T20M site on the model αGFP Fab and stability of the sulfimide linkage was measured at 37°C over 3 days. There was considerable variation in stability from 40-90% retained; nonetheless, two of oxaziridines (compound **5** and **8**) provided stability over 80% (**Fig. 2D**). In a recent parallel study, it was shown that conjugate stability to isolated methionine was related to the electron density around the carbonyl as measured by the carbonyl stretching frequency^24^. Indeed, we found a strong inverse correlation between carbonyl stretching frequency and the measured stabilities on the Fab (**Fig. 2E**) as was also seen with isolated methionine. We synthesized a new azide containing oxaziridine derivative, based on the more stable piperidine-derived oxaziridine **8**, to enable copper-free click chemistry for ADC conjugation (**Fig. 2F**).

We next investigated how lowering site accessibility may shield the resulting sulfimide linkage from hydrolysis. We knew that fully buried sites are unreactive. Therefore, we chose 23 sites that had intermediate degrees of accessibility (**Fig. 3A**, **Supplementary Table 2)** most of which were located on structured β-sheet regions. Remarkably, 19 of the 23 single methionine substitutions at these partially buried sites expressed at high levels in *E. coli* (3-50mg/L); 18 retained high affinity to GFP, and 17 retained high thermostability (**Fig. 3B**). These less accessible sites were also less reactive, and thus we increased the labeling reaction to 20 equivalents of oxaziridine to better drive the reactivity. We found four mutants that had greater than 85% stability when labeled with the oxaziridine azide **8** and incubated at 37°C for 3 days (**Fig. 3B**, **Fig. 3C**). There was a slight inverse correlation between site accessibility and long-term stability (**Fig. 3D**) but the lack of a strong correlation suggests that additional factors are at play besides simple site accessibility. Overall, we found the combination of probing different oxaziridine derivatives and different site accessibility produced highly stable conjugates that were good candidates for ADC production.

**Figure 3.**
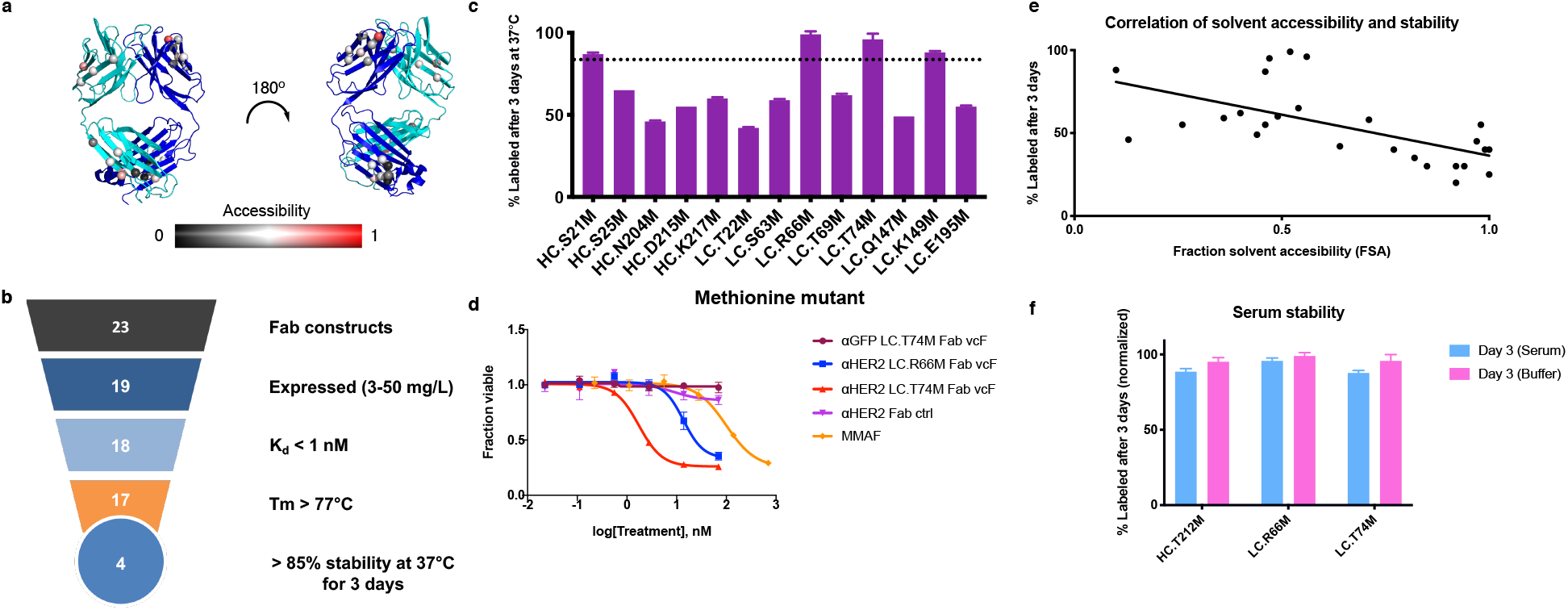
Labeling, stability, and activity of partially buried sites on αGFP-Fab and Trastuzumab Fab. (a) Structure of the trastuzumab Fab showing 23 partially buried sites (spheres) chosen to mutate to methionine. Scale represents calculated relative surface accessibility from 0 (black) to 1 (red). (b) Triage of the 23 individual mutants on the αGFP-Fab after testing expression, binding to GFP, structural stability, and oxaziridine conjugate stability for 3 days at 37°C. (c) Conjugate stability of 14 sites after incubation at 37°C for 3 days (n=3). The dotted line indicates 85% stability. (d) *In vitro* potency of two stable sites on trastuzumab Fab on the BT474-M1 breast cancer cell line (n=3). (e) Correlation between measured stability at 37°C and calculated accessibility for the 23 partially buried sites (R^2^ =0.32). (f) Stability measured in human serum for the three top stable sites over 3 days at 37°C compared to stability measured in buffer (n=3).

We next incorporated these mutations into a trastuzumab Fab and tested the ADC conjugates for killing of breast cancer cell lines. However, we noticed that the wild-type trastuzumab Fab labeled 25% with the oxaziridine reagent when reacted at 20 equivalents, which was a necessary concentration of oxaziridine to label the less accessible sites (**Supplementary Fig. 3A**). We hypothesized this additional and undesirable labeling was due to labeling of the methionine at position HC.M107 in the CDR H3 of trastuzumab. Simply mutating HC.M107 to an unreactive leucine eliminated labeling at this site (**Supplementary. Fig 3A**). Furthermore, the HC.M107L mutation did not affect binding to HER2 on SKBR3 cells (**Supplementary Fig 3B).**

We chose our two most stable sites, LC.R66M and LC.T74M, and incorporated methionine into the corresponding sites on trastuzumab αHER2 Fab antibody to use in cellular toxicity and serum stability assays. Both labeled to greater than 80% when reacted with 20 equivalents of oxaziridine-azide **8** (**Supplementary Table 2**). The two stable sites were individually converted to methionines on the trastuzumab Fab scaffold and then labeled with oxaziridine azide **8**, followed by strain promoted click chemistry with DBCO-PEG4-valine-citrulline-MMAF to be used in a cellular toxicity assay. We chose to use the cathepsin B cleavable linker valine-citrulline for its improved effect over a non-cleavable linker (data not shown). We picked the microtubule inhibitor MMAF as the toxic payload due to its previously characterized strong potency in ADC formats and improved solubility compared to MMAE^25^. Both ADCs showed high potency in a HER2-positive breast cancer cell line, BT474-M1, compared to either trastuzumab alone or an αGFP Fab control (**Fig. 3E**). The ADC conjugates were 10-100-fold more potent than the free MMAF reflecting their capacity as a drug chaperone. Interestingly, the ADC derived from the LC.R66M was about 10-fold less active than LC.T74M due to a modest loss in affinity when conjugated with drug (**Supplementary Fig. 4A**). Fortunately, upon conversion to a full IgG, the loss in affinity was greatly restored due to the higher avidity of the IgG and much lower off-rates (**Supplementary Fig. 4B**). Both sites were also tested for their stability in human serum and showed similar levels compared to their stability measured in buffer (**Fig. 3F**). Thus, the two sites in the Fab arms are promising candidates for ADC formation.

### Labeling and stability at homologous sites on the Fc domain

To explore more flexibility in labeling sites for methionine antibody conjugates, we probed for suitable labeling sites on the Fc domain of the IgG. However, we found there are two endogenous methionines on the Fc (HC.M252 and HC.M428) that are surface exposed and one of which readily reacts with the oxaziridine azide **8** (**Fig. 4A**, **Supplementary Fig. 5A**). Also, it is known that these methionines sit directly at the FcRn binding site and that even oxidation at these sites can disrupt FcRn binding^26^. We found that labeling these methionines with oxaziridine ablated FcRn binding **(Supplementary Fig. 5B).** In order to preserve FcRn binding, we chose to avoid conjugating at these sites. We scrubbed these methionines by mutation to leucine and found these had little to no effect on overall protein stability or FcRn binding ability (**Supplementary Fig. 5C**). We also incorporated an N297G mutation to prevent glycosylation of the Fc to simplify our mass spectrometry analysis. We then used this triple Fc mutant as our template to search for more stable methionine conjugation sites.

**Figure 4.**
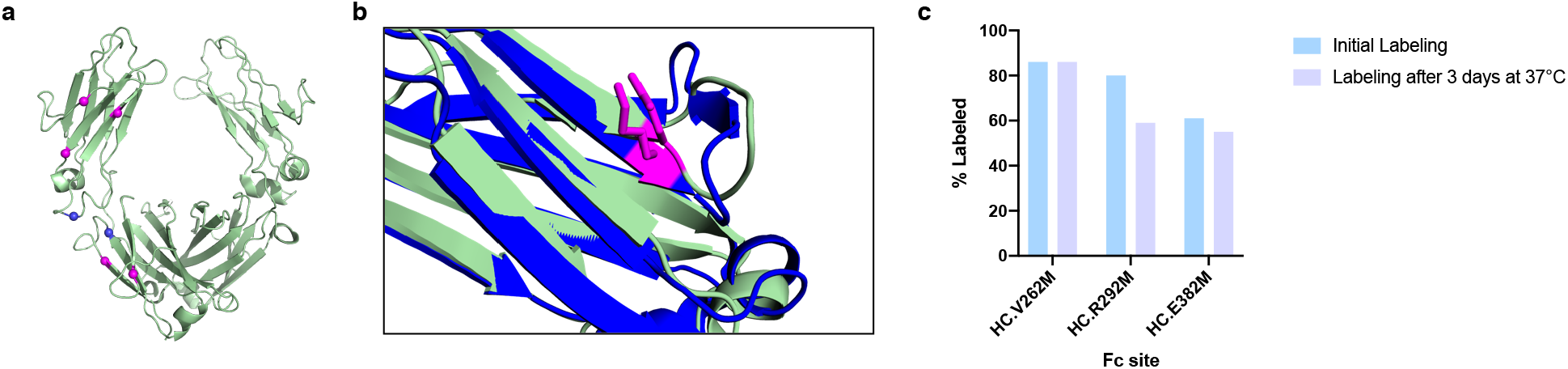
Labeling, stability, and activity of homologous Fc sites. (a) Structure of IgG1 Fc domain (PDB: 1H3X) with 5 sites chosen to individually mutate to methionine (dark purple). The two endogenous methionines are shown in magenta. (b) Example alignment of part of the Fc domain with part of the Fab light chain to show the structural homology between site LC.K149 and HC.E383. (c) Stability and labeling measurements for 3 sites on the Fc region. Two sites (HC.T307M, HC.T437M) are not shown because they did produce viable conjugates with oxaziridine.

To simplify our quest for new methionine sites in the Fc we took advantage of the high structural similarity between the Fc and Fab arms. We used PyMol to align the five most stable conjugation sites from the Fab arm studies above to sites in the Fc domain (**Fig. 4A**, **Supplementary Table 3)**. An example alignment is shown between LC.K149 and HC.E382 (Fig 4B). We introduced single methionine mutants into these sites in the native methionine-scrubbed Fc, expressed the variants in Expi293 mammalian cells, and tested them for their labeling efficiency and stability. Interestingly, two of the engineered sites (HC.T307M, HC.T437M) did not label at all and thus could not be tested for their stability. The other three sites labelled to over 50%, and site HC.V262M showed greater than 80% labeling efficiency with virtually no hydrolysis after a three day incubation at 37°C (**Fig. 4C**).

### Functional activity of methionine oxaziridine ADCs on breast cancer cell lines and in vivo efficacy in a breast cancer xenograft model

We then tested how each of the three stables sites (LC.R66M, LC.T74M, and HC.V262M) performed as ADCs in an IgG format on HER2-positive breast cancer cell lines (**Fig. 5A, 5B**). On both SKBR3 and BT474-M1 cell lines, all three sites were almost equally effective at reducing cell growth (IC_50_ ~100-1000pM). All three were 20-50-fold more potent than trastuzumab alone. When compared to one of the previously reported optimal engineered cysteine sites LC.V205C^20^, we saw comparable cell killing at sites LC.T74M and HC.V262M (**Fig. 5C**). We also tested how these conjugates performed by size exclusion chromatography (SEC) as a test for antibody aggregates and a proxy for good pharmacokinetics. ADCs produced at sites LC.T74M and HC.V262M showed a single symmetrical elution peaks comparable to trastuzumab, while site LC.R66M formed three broad peaks (**Supplementary Fig. 6**). Thus, we decided not to use site LC.R66M *in vivo*. We also discovered that after reintroducing the wildtype N297 residue and thus glycosylation, we were not able to label site HC.V262M. We hypothesized that this was because the glycans sit in the same pocket that the conjugated residue would occupy and thus the glycans prevent labeling (**Supplementary Fig. 7**). Therefore, we nominated LC.T74M as our lead candidate for *in vivo* studies. We conjugated trastuzumab IgG to valine-citrulline cleavable MMAF and performed a dose-response study in a mouse xenograft BT474-M1 breast cancer model (**Fig. 5D, 5E**). We saw dose-response efficacy and with the highest dose of 6mg/kg saw inhibition of tumor growth compared to control across 5 weeks. At 6mg/kg we also saw increased efficacy compared to trastuzumab alone, where one mouse did not respond at all to trastuzumab compared to all three mice responding to the ADC.

**Figure 5.**
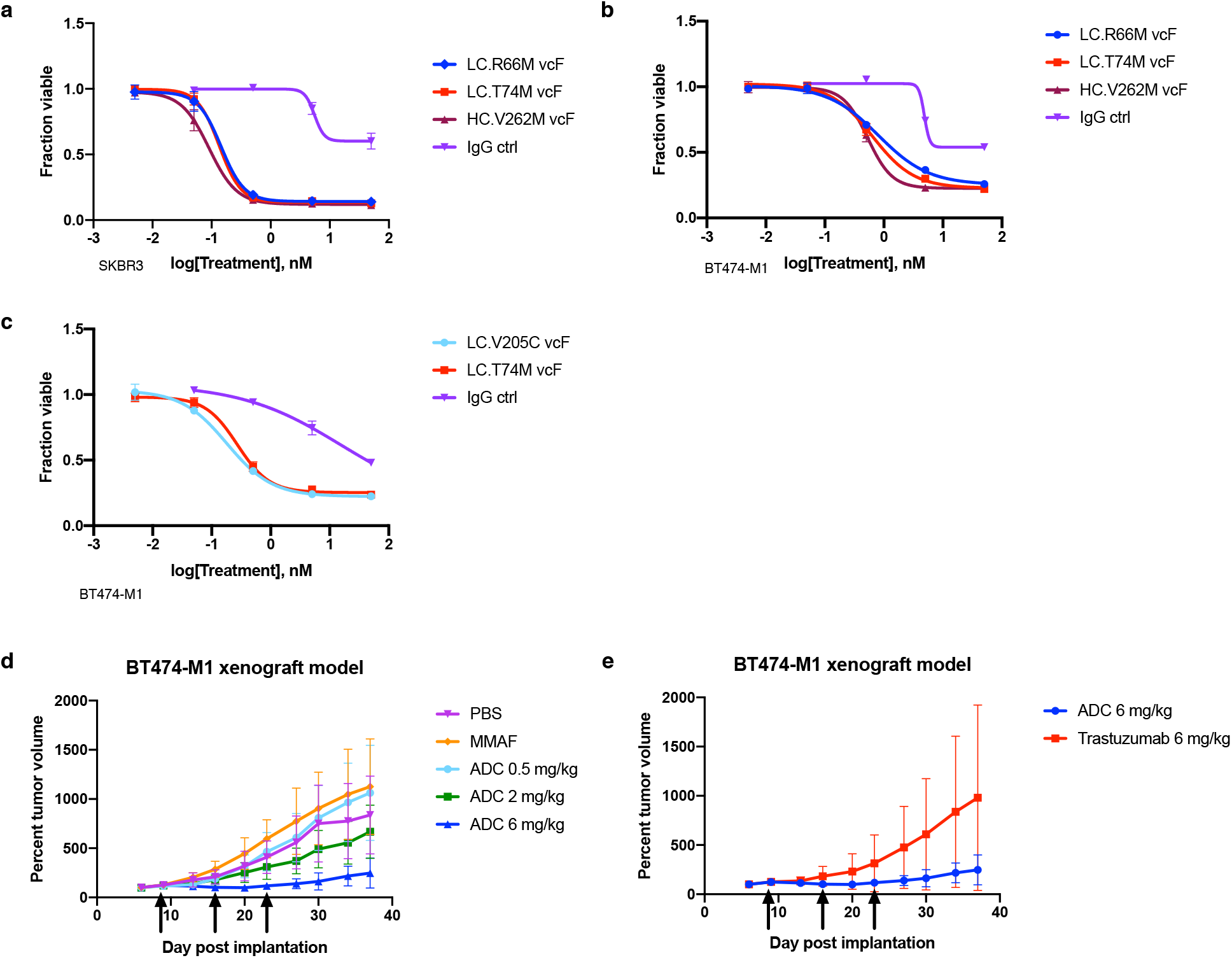
*In vitro* and *in vivo* potencies of IgG based ADCs in a breast cancer model. (a,b) *In vitro* potency of three sites (LC.R66M, LC.T74M, and HC.V262M) on two HER2 positive breast cancer cell lines, SKBR3 (a) and BT474-M1 (b) (n=3). (c) Comparison of site LC.T74M and stable cysteine site LC.V205C^20^ (n=3). (d,e) *In vivo* potency of site LC.T74M ADC in a breast cancer xenograft model in nude female mice, where (d) demonstrates clear dose response of ADC, and (e) shows the improved response to the ADC compared to the trastuzumab antibody alone (3 mice per group). Arrows show intravenous administration schemes of controls and ADCs.

## Discussion

We layout a systematic and general approach for identifying efficient, stoichiometric, and stable methionine labeling sites for antibodies using ReACT that preserves antibody function and stability for ADC applications. We explored a number of variables and addressed potential pitfalls to find optimal labeling sites. Surprisingly, almost all of the single methionine mutants were tolerated in the context of the trastuzumab scaffold. Of the 95 highly accessible methionine sites, 93 were expressed at wild-type levels and 92 retained a Tm greater than 77°C. Even for the 23 partially buried sites, 19 were expressed at wild-type levels and 17 maintained a high Tm. We did not detect methionine oxidation for the purified recombinant antibodies expressed either as Fabs in *E.coli* or as IgGs expressed from mammalian cells. This obviated the need to chemically reduce prior to conjugation with oxaziridine. This is a substantial advantage to cysteine labeling which typically requires reduction and reoxidation prior to thiol-conjugation. The conjugation to the oxaziridine was done rapidly (30 min at 5-30 fold excess) at room temperature in aqueous conditions and consistently produced high yields of the bioconjugate. For example, of the 92 accessible methionine sites expressed, 57 were labelled over 90%. Even for the 23 expressible partially buried methionine sites, 11 were labeled to over 80%.

One can tolerate, manage, or exploit endogenous methionines for antibody conjugations. In our αGFP trastuzumab Fab there are three buried methionines. We found them to be unreactive and thus preserved them throughout our experiments. Once we switched to the wild type trastuzumab there was a reactive methionine in CDR H3. This was replaced with a leucine and did not affect the affinity of the antibody. Moreover, methionines are routinely mutated out of CDRs in therapeutic antibodies to avoid oxidation upon long-term storage or treatment^27^. We also identified two endogenous methionines in the Fc and these were readily mutated to leucine without significant impact on expression or binding. In fact, in some cases these sites have been mutated away from methionine to extend antibody half-life and improve FcRn binding^28^.

We found the initial oxaziradine compound did not have the desired stability for long-term studies, but structure-activity analysis identified new compounds with significantly improved stability to hydrolysis. The stability tracked with the electron density surrounding the carbonyl as found in parallel studies on isolated methionine^29^. We believe these new compounds (especially oxaziridine azide **8**) will find general utility for ReACT applications for other protein bioconjugations.

We found significant variation in stability depending on the site of modification. There is an inverse trend between accessibility and site stability. We expect this may be because the sulfimide is shielded from water and hindered from being hydrolyzed. The hydrolysis reaction of the sulfimide is expected to go through a tetrasubstituted intermediate^30^ and neighboring sites will likely impact the stability of this intermediate based on the chemical environment. Further mechanistic and computational work will help to further dissect these factors. Interestingly, the stability, and therefore therapeutic effectivity, of cysteine conjugates also varies depending on the conjugation sites^19,31^. While we see that very accessible sites tend to be reactive, there is no clear trend between accessibility and reactivity, as was also seen with cysteine sites^19^.

We believe site-specific modification of methionine by ReACT has great potential for antibody and protein bioconjugations. The expression of the methionine mutants is robust and the general tolerability of methionine mutations suggests multiple methionines could easily be introduced. The conjugation procedure is rapid, simple, and does not require pre-reduction. There is good flexibility with site selection and the resulting linkage can be stable at biological temperatures. The sites described here will provide candidates for other antibody scaffolds. In fact, the discovery of the stable Fc site did not require a complete methionine surface scan, but rather simple homology modeling was sufficient to identify useful sites. Site-specific methionine labeling by ReACT offers more homogeneity of modification compared to lysine modification. It produced conjugates as stable as cysteine-maleimide conjugations, and robust ADC activity in a BT474-M1 mouse xenograft model. While there is still much to do to validate their clinical use, the methionine modification path looks promising. This modification will be useful for many other antibody and protein bioconjugation applications such as for fluorescence, affinity labels, DNA barcoding, and protein-protein bioconjugation. We believe the general parameters we analyze and optimize here will expand the use of ReACT bioconjugation on many other biomolecules.

## Abbreviations

ADC: Antibody drug conjugate
DAR: drug-antibody-ratio
ReACT: Redox-activated chemical tagging
vcF: DBCO-PEG4-valine-citrulline-MMAF
CDR: Complementarity determining region
DSF: Differential scanning fluorimetry

## Acknowledgements

We thank the members of the Wells laboratory and Antibiome for helpful discussions. We thank M. Hornsby for the αGFP Fab expression vector, A. Weeks for the αHER2 Fab expression vector, A. Cotton for the V205C mutant vector, and J. Zhou for input on the cell viability assay. J.A.W. thanks The Chan Zuckerberg Initiative and Biohub Investigator Program as well as NCI grant P41CA196276 for financial support of this work. H.L.T. was supported from NIH R21 AI111662. S.K.E. thanks the NSF GRFP (DGE 1650113) for financial support. F.D.T. thanks Novartis Institutes for BioMedical Research and the Novartis-Berkeley Center for Proteomics and Chemistry Technologies (NB-CPACT) for supporting this work. A.H.C. thanks the NSF-GRFP (DGE 1106400) for financial support. C.J.C. acknowledges the NIH (ES4705 and ES28096) and the Aduro-Berkeley IVRI program for financial support. C.J.C. is an Investigator with the Howard Hughes Medical Institute and a CIFAR Senior Fellow.

## Author contributions

S.K.E., H.L.T., and J.A.W. designed the research. S.K.E and H.L.T. performed the mutant work, and S.K.E. performed the compound screen, stability assays, and ADC assays. A.H.C synthesized the oxaziridine compounds and performed the stretching frequency correlation, under guidance from F.D.T and C.J.C. Animal experiments were performed by V.S., B.H., and UCSF PTC. S.K.E., H.L.T., and J.A.W. analyzed data and interpreted results. S.K.E. and J.A.W. wrote the manuscript and all provided editorial comments.

## Materials and correspondence

Correspondence and material requests should be addressed to J.A.W.

## Materials and Methods

### Selection of accessible conjugation sites

To estimate the relative solvent accessibility (RSA) of engineered methionines on a Fab, a computational methionine scan was performed with MODELLER using PDB structure 1FVE as a template^32^. MODELLER generates homology models for comparative structure analysis by satisfaction of spatial restraints^33^. Single methionine mutations were systematically modeled across the entire structure of the Fab including an additional model with a methionine appended at the end of the light chain for a total of 439 individual models generated. The solvent accessible surface area (SASA) of the engineered methionine sulfur atom was determined using the “get_area” function (dot_solvent = 1, dot_density = 4, solvent_radius = 1.4) in PyMol. Due to the stochasticity of the S-methyl group placement, the group was removed prior to SASA calculations and was found to reduce variability. The RSA was calculated by taking the SASA values and dividing by the maximum SASA value observed in the set. Positions were rank ordered and the top 95 sites with the highest RSA (excluding CDR positions, prolines and cysteines) were selected for bioconjugation.

### Preparation and characterization of αGFP Fab methionine mutants

All methionine mutants were made using QuikChange to introduce single codon mutations onto the αGFP Fab. Fabs were expressed and purified by an optimized auto-induction protocol previously described^23^. In brief, C43 (DE3) Pro +*E. coli* containing expression plasmids were grown in TB auto-induction media at 37 °C for 6 hours, then cooled to 30 °C for 16–18 hr. Cells were harvested by centrifugation and Fabs were purified by Protein A affinity chromatography. Fab purity and integrity was assessed by SDS-PAGE and intact protein mass spectrometry using a Xevo G2-XS Mass Spectrometer (Waters) equipped with a LockSpray (ESI) source and Acquity Protein BEH C4 column (2.1 mm inner diameter, 50 mm length, 300 Å pore size, 1.7 μm particle size) connected to an Acquity I-class liquid chromatography system (Waters). Deconvolution of mass spectra was performed using the maximum entropy (MaxEnt) algorithm in MassLynx 4.1 (Waters).

### Labeling of αGFP Fab methionine mutants with oxaziridine and Sulfo-DBCO-NHS

Fabs were prepared at 30uM in PBS and labeled with 5 equivalents of the original oxaziridine azide reagent. The reaction proceeded for 2 hours at room temperature before being quenched with 500mM methionine. Sulfo-DBCO-NHS was added at a final concentration of 625uM and incubated at room temperature. Labeling was analyzed by intact protein mass spectrometry using a Xevo G2-XS Mass Spectrometer as previously described.

### Single-point kinetic screen

To determine if binding was perturbed by conjugation, a single-point kinetic screen was performed by bio-layer interferometry on a ForteBio Octet RED384. Biotinylated-GFP was captured by streptavidin biosensors and the remaining biotin binding sites were saturated with free biotin. Association of 10 nM unlabeled or labeled Fab was measured for 15 min followed by dissociation for 30 min. K_D_ values of all unlabeled and labelled Fabs were estimated to be sub- 0.5nM. Binding affinity for FcRn was performed in a similar manner but at pH 6.0 to mimic binding in the acidic endosome. Biotinylated FcRn (Acro Biosystems) was used as the loading ligand.

### Protein stability Differential Scanning Fluorimetry (DSF) assay

Stability was measured by a Sypro Orange based DSF assay. In brief, Fabs (2μM) were incubated with 4x Sypro Orange Protein Gel Stain (ThermoFischer) in PBS. Fluorescence scanning was performed from 25°C- 95°C at a rate of 1°C/min using a Lightcycler 480 Instrument (Roche Life Scientific). Melting temperatures were calculated from the inflection point in the first-derivative curve.

### Synthesis of Compounds

All oxaziridines compounds were previously synthesized and reported in Christian *et al*^29^. Synthesis of the azide-piperdine oxaziridine (oxaziridine azide **8**) can be found in the supplementary methods.

### Parameter Derivation

A conformational search on the respective ureas and carbamates was performed using the MacroModel suite from Schrödinger^34^ using an OPLS_2005 force field without solvent corrections. A Monte-Carlo molecular mechanics method was employed. The output was restricted to structures within 1.30 kcal/mol (5 kJ/mol) of the lowest energy conformer. Conformers were submitted to a geometry optimization in Gaussian 09 using the def2-TZVP basis set and M06-2x functional^35^. A triple zeta potential basis set was chosen along with the M06-2x functional, as these generally lead to quantitative correlations^36^. Using a cutoff limit of 2.5 kcal/mol, the parameters of each low energy conformer were weighted using the Boltzmann distribution (equations 1 and 2) where the energy of a given conformer is calculated relative to the lowest energy conformation.

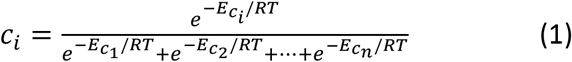

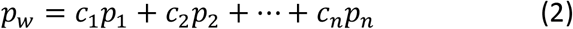

### PyMol homology alignment

To determine analogous stable sites on the Fc, the alignment function was used in PyMol, using the PDB structure 1FVE (Fab) and 1H3X (Fc). Stable sites on light chain or heavy chain were aligned to either the CH2 or CH3 domains in the Fc. Corresponding positions were chosen on the Fc to mutate to methionine.

### Expression of IgG single methionine mutants

IgGs containing the engineered methionines were expressed and purified from Expi293 BirA cells according to established protocol from the manufacturer. Briefly, 30μg of pFUSE (InvivoGen) vector was transiently transfected into 75 million Expi293 BirA cells using the Expifectamine kit. Enhancer was added 20 hours after transfection. Cells were incubated for a total of 6 days at 37 °C in a 5% CO_2_ environment before the supernatants were harvested by centrifugation. Protein was purified by Protein A affinity chromatography and assessed for quality and integrity by SDS-PAGE.

### Conjugation of engineered methionine Fabs and IgGs with oxaziridine and DBCO-PEG4-valcit-MMAF

For Fab ADCs, endotoxins were removed prior to conjugation using Pierce endotoxin removal kits (ThermoFischer Scientific). For conjugation, Fabs were incubated at 50μM with 15 molar equivalents of compound 8 azide oxaziridine for 30 minutes at room temperature in PBS. For IgGs, IgGs were incubated at 10μM with 30 molar equivalents of compound 8 azide oxaziridine per methionine for 1 hour at room temperature in PBS. For both, the reaction was quenched by the addition of methionine and antibody was buffered exchanged into PBS using a 0.5mL Zeba 7kDa desalting column (ThermoFischer Scientific). Then 10 molar equivalents of DBCO-PEG4-valcit-MMAF (Levena Biosciences) was added and the click reaction proceeded overnight at room temperature. The conjugate was desalted twice using two 0.5mL Zeba 7kDa columns to remove excess unconjugated drug. Full conjugation was monitored by intact protein mass spectrometry using a Xevo G2-XS Mass Spectrometer (Waters).

### Conjugation of engineered cysteine ADCs for comparison

Engineered cysteine conjugation was performed as previously reported^37^. In brief, after purification of the LC.V205C mutant αHer2 IgG (see IgG expression), the IgG (10μM) was buffer exchanged into 50mM Tris-HCl, pH 7.5, 2mM EDTA. DTT was added at 40-fold molar excess and incubated at room temperature for 16 hours. Desalting into PBS proceeded with 0.5mL Zeba 7kDa columns. DHAA was added in 15-fold molar excess to reoxidize the interchain disulfides for 3 hours at room temperature. Maleimide-valcit-MMAF (BOC Sciences) was added at 3-fold molar excess and conjugation was monitored by mass spectrometry. Excess drug was removed by two 0.5mL Zeba desalting columns.

### Cell culture of HER2-positive breast cancer cells

The BT474-M1 cell line was provide by the Preclinical Therapeutics Core at the UCSF Helen Diller Cancer Center. These cells were maintained in DMEM media supplemented 10% FBS and 1X Pen/Strep. The SKBR3 cells were purchased from the UCSF Cell Culture Facility. They were maintained in McCoy 5a media supplemented with 10% FBS and 1X Pen/Strep. Cell line identities were authenticated by morphological inspection. The SKBR3 cell line identity was validated by UCSF Cell Culture Facility. Symptoms for mycoplasma contamination were not observed and thus no test for mycoplasma contamination was performed. All cell lines that were received as gifts were previously authenticated and tested for mycoplasma.

### ADC cell killing assay in vitro

Antibody drug conjugate cell killing assays were performed using an MTT modified assay to measure cell viability. In brief, 10000 BT474-M1 or SKBR3 cells were plated in each well of a 96-well plate on day 0. On day 1, Fab/IgG was added in a 10-fold dilution series. Cells were incubated for 120 hr at 37°C under 5% CO_2_. On day 6, 40uL of 2.5mg/mL of Thiazolyl Blue Tetrazolium Bromide (Sigma Aldrich) was added to each well and incubated at 37°C under 5% CO_2_ for 4 hours. Following, 100μL of 10% SDS 0.01M HCl was added to lyse the cells to release the MTT product. After 4 hours, absorbance at 600nm was quantified using an Infinite M200 PRO plate reader (Tecan).

### ADC study in mouse xenograft model in vivo

The xenograft was performed with 6-8 week old nude female mice (NCR, nu/nu) purchased from Taconic Labs (n=3 per group). Prior to tumor cell engraftment, mice were implanted subcutaneously with Estradiol pellet (0.36mg, 60 day release, Innovative Research). BT474-M1 xenografts were then established by bilateral subcutaneous injection into the right and left flanks of mice with BT474-M1 tumor cells (5×10^6^ cells in 100 μl of serum free medium mixed 1:1 with Matrigel). When BT474-M1 xenografts reached average volume of 200mm^3^ (measured as width × width × length × 0.52), mice were dosed intravenously weekly for 3 weeks with PBS, drug alone, antibody alone and ADCs. Tumor size and body weight were monitored biweekly for 5 weeks total.

## References

1. Adams, G. P. & Weiner, L. M. Monoclonal antibody therapy of cancer. Nature Biotechnology (2005). doi:10.1038/nbt1137

2. Mao, S.-Y. & Mullins, J. M. Conjugation of fluorochromes to antibodies. Methods Mol. Biol. 588, 43–8 (2010).

3. Meares, C. F. et al. Conjugation of antibodies with bifunctional chelating agents: isothiocyanate and bromoacetamide reagents, methods of analysis, and subsequent addition of metal ions. Anal. Biochem. 142, 68–78 (1984).

4. Darmanis, S. et al. Simultaneous Multiplexed Measurement of RNA and Proteins in Single Cells. Cell Rep. 14, 380–389 (2016).

5. Stoeckius, M. et al. Large-scale simultaneous measurement of epitopes and transcriptomes in single cells. (2017).

6. Tushir-Singh, J. Antibody-siRNA conjugates: drugging the undruggable for anti-leukemic therapy. Expert Opin. Biol. Ther. 17, 325–338 (2017).

7. Lewis Phillips, G. D. et al. Targeting HER2-positive breast cancer with trastuzumab-DM1, an antibody-cytotoxic drug conjugate. Cancer Res. 68, 9280–90 (2008).

8. Lynch, C. M. et al. Brentuximab Vedotin (SGN-35) for Relapsed CD30-Positive Lymphomas. N. Engl. J. Med. (2010). doi:10.1056/nejmoa1002965

9. DiJoseph, J. F. et al. Antibody-targeted chemotherapy with CMC-544: a CD22-targeted immunoconjugate of calicheamicin for the treatment of B-lymphoid malignancies. Blood 103, 1807–14 (2004).

10. Egan, P. C. & Reagan, J. L. The return of gemtuzumab ozogamicin: a humanized anti-CD33 monoclonal antibody-drug conjugate for the treatment of newly diagnosed acute myeloid leukemia. Onco. Targets. Ther. 11, 8265–8272 (2018).

11. Beck, A., Goetsch, L., Dumontet, C. & Corvaïa, N. Strategies and challenges for the next generation of antibody-drug conjugates. Nature Reviews Drug Discovery (2017). doi:10.1038/nrd.2016.268

12. Panowski, S., Bhakta, S., Raab, H., Polakis, P. & Junutula, J. R. Site-specific antibody drug conjugates for cancer therapy. MAbs 6, 34–45 (2014).

13. Chalouni, C. & Doll, S. Fate of Antibody-Drug Conjugates in Cancer Cells. J. Exp. Clin. Cancer Res. 37, 1–12 (2018).

14. Tsuchikama, K. & An, Z. Antibody-drug conjugates: recent advances in conjugation and linker chemistries. Protein and Cell (2018). doi:10.1007/s13238-016-0323-0

15. Hamblett, K. J. et al. Effects of Drug Loading on the Antitumor Activity of a Monoclonal Antibody Drug Conjugate. 10, 1–9 (2013).

16. Bryant, P. et al. In vitro and in vivo evaluation of cysteine rebridged trastuzumab-MMAE antibody drug conjugates with defined drug-to-antibody ratios. Mol. Pharm. 12, 1872–1879 (2015).

17. Liu-Shin, L. P.-Y., Fung, A., Malhotra, A. & Ratnaswamy, G. Evidence of disulfide bond scrambling during production of an antibody-drug conjugate. MAbs 10, 1190–1199

18. Junutula, J. R. et al. Rapid identification of reactive cysteine residues for site-specific labeling of antibody-Fabs. J. Immunol. Methods 332, 41–52 (2008).

19. Ohri, R. et al. High-Throughput Cysteine Scanning to Identify Stable Antibody Conjugation Sites for Maleimide- and Disulfide-Based Linkers. Bioconjug. Chem. 29, 473–485 (2018).

20. Lin, S. et al. Redox-based reagents for chemoselective methionine bioconjugation. Science (80-.). 602, 597–602 (2017).

21. Dyer, K. F. The Quiet Revolution: A New Synthesis of Biological Knowledge. J. Biol. Educ. (1971). doi:10.1080/00219266.1971.9653663

22. Persson, H. et al. CDR-H3 diversity is not required for antigen recognition by synthetic antibodies. J. Mol. Biol. 425, 803–811 (2013).

23. Hornsby, M. et al. A High Through-put Platform for Recombinant Antibodies to Folded Proteins. Mol. Cell. Proteomics 14, 2833–47 (2015).

24. Christian, A. H. et al. A Physical Organic Approach to Tuning Reagents for Selective and Stable Methionine Bioconjugation. J. Am. Chem. Soc. jacs.9b04744 (2019). doi:10.1021/jacs.9b04744

25. Mendelsohn, B. A. et al. Investigation of Hydrophilic Auristatin Derivatives for Use in Antibody Drug Conjugates. Bioconjug. Chem. 28, 371–381 (2017).

26. Gao, X. et al. Effect of individual Fc methionine oxidation on FcRn binding: Met252 oxidation impairs FcRn binding more profoundly than Met428 oxidation. J. Pharm. Sci. 104, 368–77 (2015).

27. Haberger, M. et al. Assessment of chemical modifications of sites in the CDRs of recombinant antibodies: Susceptibility vs. functionality of critical quality attributes. MAbs (2014). doi:10.4161/mabs.27876

28. Kuo, T. T. & Aveson, V. G. Neonatal Fc receptor and IgG-based therapeutics. mAbs (2011). doi:10.4161/mabs.3.5.16983

29. Christian, A. H. et al. A Physical Organic Approach to Tuning Reagents for Selective and Stable Methionine Bioconjugation. ChemRxiv (2019).

30. Pyun, S. Y., Kim, T. R., Lee, C. R. & Kim, W. G. Kinetics studies on the mechanism of hydrolysis of S-phenyl-S-vinyl-N-p-tosylsulfilimine derivatives. Bull. Korean Chem. Soc. 24, 306–310 (2003).

31. Shen, B. Q. et al. Conjugation site modulates the in vivo stability and therapeutic activity of antibody-drug conjugates. Nat. Biotechnol. (2012). doi:10.1038/nbt.2108

32. Eigenbrot, C., Randal, M., Presta, L., Carter, P. & Kossiakoff, A. A. X-ray structures of the antigen-binding domains from three variants of humanized anti-p185HER2 antibody 4D5 and comparison with molecular modeling. J. Mol. Biol. (1993). doi:10.1006/jmbi.1993.1099

33. Webb, B. & Sali, A. Comparative protein structure modeling using MODELLER. Curr. Protoc. Bioinforma. (2016). doi:10.1002/cpbi.3

34. Schrödinger Release 2014-3: MacroModel, Schrödinger, LLC, New York, NY, 2014. No Title.

35. Gaussian 09, Revision A.02, M. J. Frisch, G. W. Trucks, H. B. Schlegel, G. E. Scuseria, M. A. Robb, J. R. Cheeseman, G. Scalmani, V. Barone, G. A. Petersson, H. Nakatsuji, X. Li, M. Caricato, A. Marenich, J. Bloino, B. G. Janesko, R. Gomperts, B. Mennu, 2016. No Title.

36. Zhao, Y. & Truhlar, D. G. The M06 suite of density functionals for main group thermochemistry, thermochemical kinetics, noncovalent interactions, excited states, and transition elements: Two new functionals and systematic testing of four M06-class functionals and 12 other functionals. Theor. Chem. Acc. (2008). doi:10.1007/s00214-007-0310-x

37. Sassoon, I. & Blanc, V. Antibody–Drug Conjugate (ADC) Clinical Pipeline: A Review. in 1–27 (2013). doi:10.1007/978-1-62703-541-5_1

