## Supplementary Information for "Systematic identification of engineered methionines and oxaziridines for efficient, stable, and site-specific antibody bioconjugation"

**Supplementary Figures/Tables:**

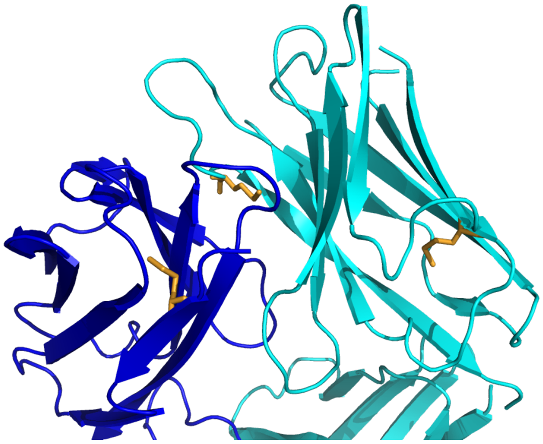

**Supplementary Fig 1. Native methionines in the trastuzumab Fab.**

Methionines present in the trastuzumab Fab are shown in orange. One methionine (LC.M04) is present in the variable domain of the light chain (dark blue) and two (HC.M86, HC.M107) are present on the heavy chain (cyan), with HC.M107 present on the H3 loop of the CDR binding domain.

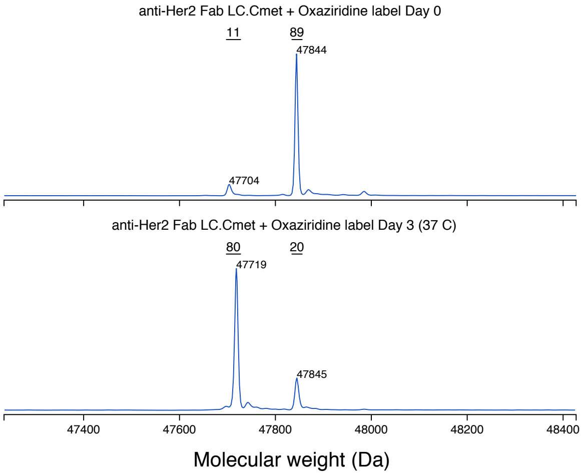


**Supplementary Fig 2. Stability of the light chain C-terminal methionine at 37°C.**

Incubation of the C-terminal methionine oxaziridine conjugate results in 80% hydrolysis over three days at the biological temperature of 37°C.

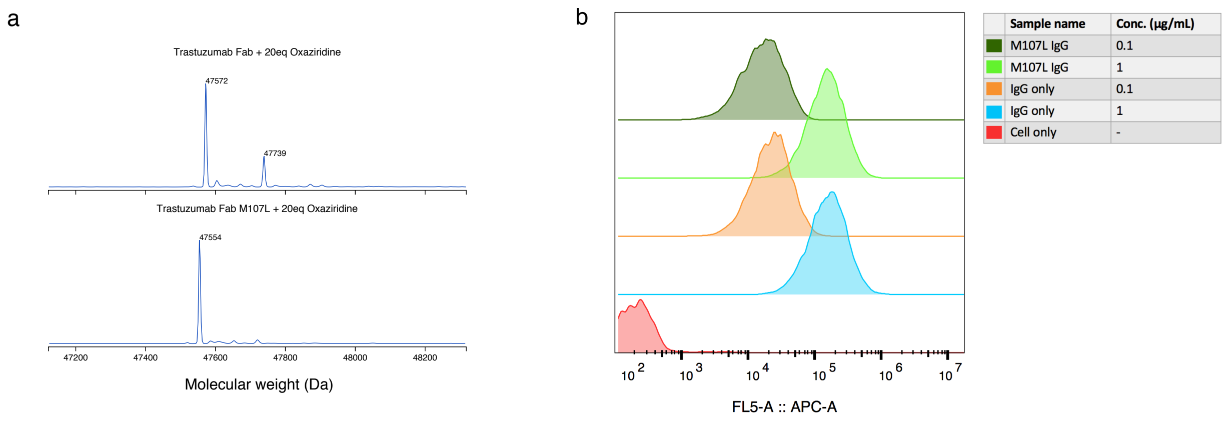


**Supplementary Fig 3. Labeling of trastuzumab Fab HC M107 and mutation to leucine prevents further labeling and minimal change in affinity for HER2.**

(a) Native trastuzumab Fab begins to be labeled with oxaziridine at equivalents of 20 or higher. However, upon mutation of M107 to leucine, this labeling is abolished.

(b) When in the IgG format, there is no detectable difference in affinity for HER2 between trastuzumab and M107L trastuzumab mutant, as detected by flow cytometry on SKBR3 cells.

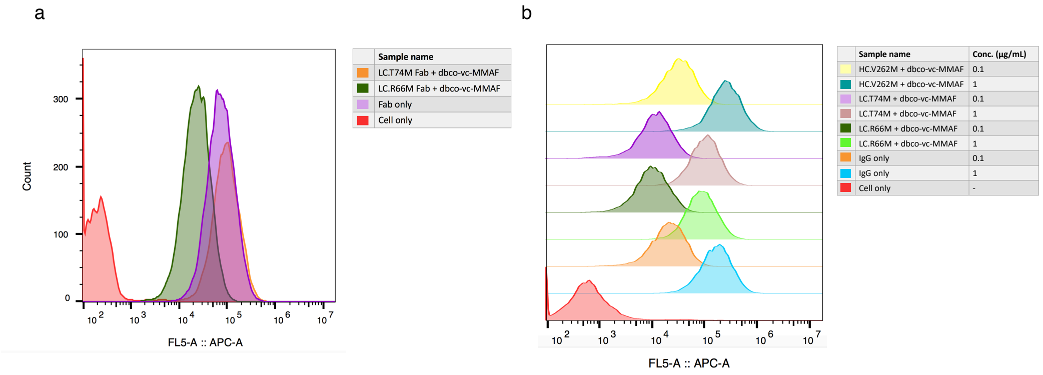


**Supplementary Fig 4. Affinity for HER2 of antibody drug conjugates in Fab and IgG format.**

(a) Flow cytometry was performed on BT474-M1 cells to confirm binding of Fab-drug conjugates at different methionine sites. LC.T74M shows similar affinity to HER2 as the unmodified trastuzumab Fab, while LC.R66M takes about a 10-fold hit in affinity. (b) Upon conversion to IgG and labeling with drug, both sites along with HC.V262M show high affinity for HER2 at two different concentrations, as determined by flow cytometry. All have roughly similar binding shifts as unlabeled IgG.

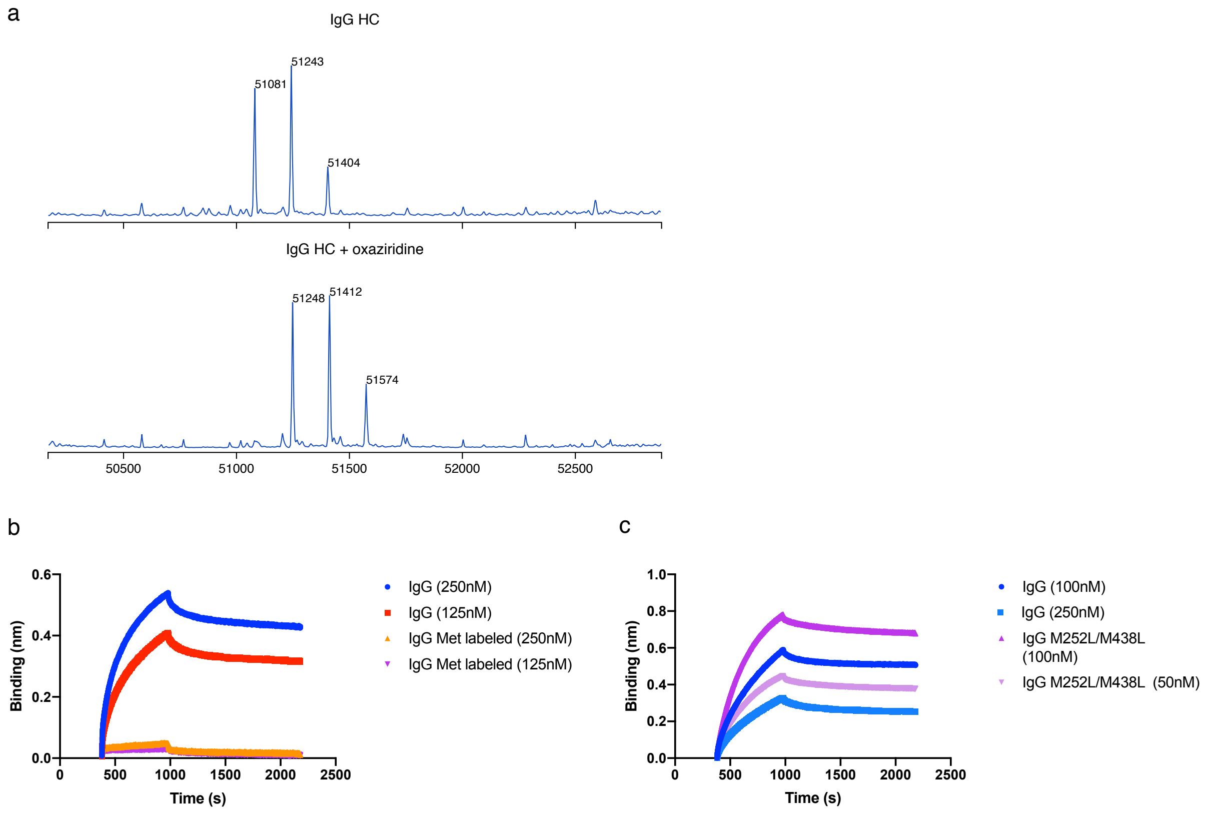


**Supplementary Fig 5. Stability of endogenous Fc methionines and mutation to leucine does not affect FcRn binding.**

(a) Rapid labeling of one methionine on the native Fc heavy chain (HC) with 30 equivalents of oxaziridine. The three peaks are different glycosylation forms of the antibody. All peaks shift 167 demonstrating reaction with the oxaziridine azide **8**.

(b) BLI analysis demonstrates that Fc methionines have undetectable binding to FcRn after being labeled with oxaziridine (orange and purple), while unlabeled IgG shows good binding to FcRn (red and blue).

(c) BLI analysis demonstrates a very similar binding affinity for FcRn of the native Fc of the IgG (blue) and M252L/M438L Fc of the IgG (purple), as noted by the similar shift by octet.

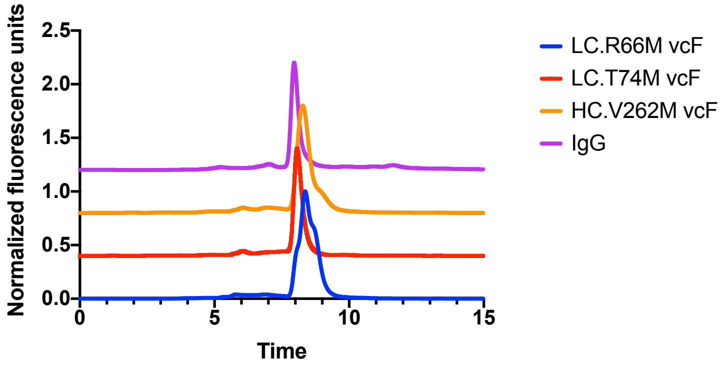


**Supplementary Fig 6. Size exclusion chromatography (SEC) of drug labeled IgG ADCs**.

SEC analysis across three sites of drug labeled IgGs compared to an unlabeled trastuzumab IgG. Both drug labeling at sites LC.T74M and HC.V262M show similar SEC profiles to unlabeled IgG, but site LC.R66M shows a drastically different SEC profile shape.

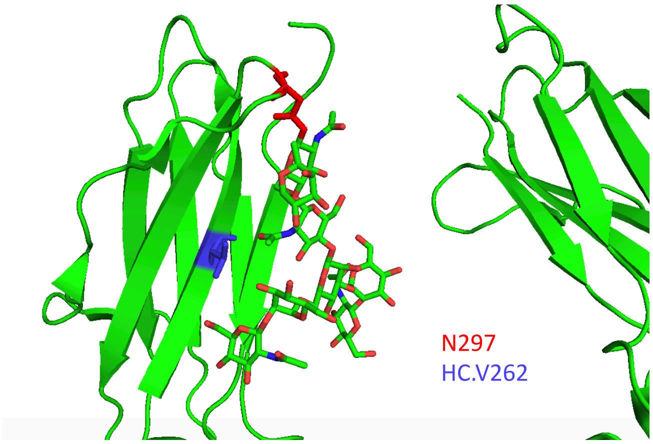


**Supplementary Fig 7. Crystal structure showing the position of Fc glycosylation and site HC.V262**.

Crystal structure of a glycosylated Fc (PDB: 5XJF) demonstrates that the HC.V262 site (shown in purple) sits right beneath the glycans. Thus, it is likely that presence of these glycans inhibits oxaziridine conjugation to HC.V262M.

**Table 1: Labeling, stability, and affinity of 95 methionine substitutions in model αGFP-Fab (rAB1001) in trastuzumab scaffold.**

| | Residue | Kabat | ASA | Yield (mg/L) | % Labeled | $K_d$ (pM) | | Thermostability (°C) | |
|---|---|---|---|---|---|---|---|---|---|
| | | | | | | Pre | Post | Pre | Post |
| rAB1001 | - | - | 0 | 8 | 0 | 88 | 197 | 82.5 | 82.6 |
| LC-Cmet | - | - | 0.67 | 7.2 | 93 | 129 | 183 | 82.4 | 82.2 |
| LC001 | D | 1 | 0.94 | 7.1 | 76 | 118 | 293 | 83.2 | 82.5 |
| LC003 | Q | 3 | 0.95 | 0 | 85 | 138 | 58 | 82.5 | 82.2 |
| LC007 | S | 7 | 0.98 | 5.4 | 96 | 111 | 150 | 82.6 | 81.5 |
| LC009 | S | 9 | 0.86 | 16.9 | 93 | 96 | 339 | 81.9 | 81.5 |
| LC010 | S | 10 | 0.73 | 7.2 | 94 | 50 | 189 | 82.9 | 82.2 |
| LC012 | S | 12 | 0.67 | 8.3 | 63 | 115 | 169 | 81.7 | 80.3 |
| LC014 | S | 14 | 0.9 | 6.4 | 93 | 112 | 162 | 82.3 | 81.8 |
| LC018 | R | 18 | 0.86 | 10.9 | 80 | 80 | 246 | 82 | 81.2 |
| LC020 | T | 20 | 0.71 | 9.6 | 95 | 80 | 183 | 81.8 | 80.6 |
| LC041 | G | 41 | 0.92 | 7.3 | 95 | 132 | 175 | 81.7 | 82.4 |
| LC042 | K | 42 | 0.79 | 7.8 | 87 | 101 | 155 | 81.8 | 81.2 |
| LC045 | K | 45 | 0.73 | 5.9 | 93 | 167 | 216 | 80.9 | 79.9 |
| LC057 | G | 57 | 0.94 | 9.1 | 93 | 123 | 252 | 80.3 | 79.9 |
| LC060 | S | 60 | 1 | 6.3 | 92 | 125 | 267 | 81.7 | 81 |
| LC065 | S | 65 | 0.9 | 11.8 | 92 | 82 | 285 | 81.6 | 81.5 |
| LC067 | S | 67 | 1 | 6.6 | 94 | 120 | 205 | 82.2 | 81.9 |
| LC068 | G | 68 | 0.9 | 6.5 | 91 | 88 | 191 | 80.2 | 79.6 |
| LC069 | T | 69 | 0.83 | 11.4 | 69 | 103 | 170 | 82.3 | 81.7 |
| LC076 | S | 76 | 0.74 | 6.3 | 82 | 135 | 175 | 81.9 | 80.7 |
| LC081 | E | 81 | 0.69 | 6.4 | 84 | 116 | 213 | 79.9 | 78.8 |
| LC085 | T | 85 | 0.68 | 8.3 | 61 | 118 | 147 | 81.6 | 80.2 |
| LC100 | Q | 100 | 0.86 | 5.2 | 90 | 116 | 74 | 81.1 | 80.9 |
| LC107 | K | 107 | 0.84 | 5.1 | 90 | 148 | 149 | 81.5 | 80.1 |
| LC108 | R | 108 | 0.8 | 0 | - | - | - | - | - |
| LC112 | A | 112 | 0.73 | 6.4 | 88 | 94 | 201 | 80.2 | 77.7 |
| LC114 | S | 114 | 0.83 | 4.7 | 89 | 165 | 207 | 82.5 | 81.3 |
| LC123 | E | 123 | 0.76 | 8.1 | 57 | 147 | 183 | 82.3 | 81.8 |

| | | | | | | | | | |
|---|---|---|---|---|---|---|---|---|---|
| LC126 | K | 126 | 0.87 | 7.7 | 87 | 97 | 198 | 82.3 | 82.3 |
| LC128 | G | 128 | 0.98 | 6.5 | 90 | 94 | 177 | 81.6 | 81.3 |
| LC143 | E | 143 | 0.94 | 7.6 | 87 | 129 | 138 | 81.8 | 81.5 |
| LC145 | K | 145 | 0.82 | 6.3 | 93 | 93 | 190 | 82.4 | 82.2 |
| LC152 | N | 152 | 0.71 | 4.3 | 88 | 95 | 192 | 82 | 82.1 |
| LC156 | S | 156 | 0.99 | 9.8 | 89 | 124 | 190 | 82.6 | 82.3 |
| LC157 | G | 157 | 1 | 7.4 | 86 | 116 | 223 | 82 | 82 |
| LC160 | Q | 160 | 0.77 | 4.8 | 58 | 94 | 169 | 82.7 | 80.6 |
| LC168 | S | 168 | 0.92 | 0 | - | - | - | - | - |
| LC169 | K | 169 | 1 | 5.1 | 92 | 264 | 179 | 82 | 81.9 |
| LC182 | S | 182 | 0.9 | 8.1 | 89 | 92 | 168 | 82.3 | 81.6 |
| LC184 | A | 184 | 0.94 | 3.6 | 71 | 110 | 148 | 82.3 | 82.2 |
| LC190 | K | 190 | 1 | 4.4 | 94 | 227 | 202 | 82.5 | 82.3 |
| LC202 | S | 202 | 0.92 | 6.8 | 93 | 103 | 237 | 82.3 | 82.2 |
| LC203 | S | 203 | 1 | 6.3 | 92 | 211 | 152 | 82.3 | 82.2 |
| LC205 | V | 205 | 0.76 | 7.7 | 71 | 97 | 190 | 82.1 | 80.8 |
| LC210 | N | 210 | 0.72 | 7.6 | 88 | 79 | 178 | 82.6 | 82.4 |
| HC001 | E | 1 | 0.78 | 6.3 | 92 | 101 | 141 | 82.3 | 82.2 |
| HC003 | Q | 4 | 0.85 | 4.5 | 94 | 94 | 149 | 82.5 | 82.6 |
| HC005 | V | 5 | 0.89 | 4.9 | 96 | 139 | 111 | 82.4 | 82.5 |
| HC007 | S | 7 | 0.95 | 7.4 | 95 | 128 | 156 | 82.3 | 82.6 |
| HC008 | G | 8 | 0.72 | 5.2 | 95 | 141 | 189 | 81.2 | 81 |
| HC010 | G | 10 | 0.97 | 7.2 | 85 | 113 | 219 | 81.5 | 81.1 |
| HC013 | Q | 13 | 0.69 | 4.1 | 89 | 133 | 138 | 82.1 | 81.7 |
| HC015 | G | 15 | 0.98 | 4.9 | 92 | 114 | 170 | 81.8 | 81.6 |
| HC017 | S | 17 | 0.75 | 5.2 | 96 | 84 | 137 | 81.9 | 82 |
| HC019 | R | 19 | 0.86 | 4.3 | 88 | 83 | 47 | 82.3 | 81.9 |
| HC023 | A | 23 | 0.86 | 5.2 | 94 | 54 | 153 | 82.6 | 82.6 |
| HC042 | G | 42 | 1 | 5.4 | 96 | 143 | 183 | 81.8 | 82.2 |
| HC043 | K | 43 | 0.89 | 6.3 | 87 | 122 | 122 | 82.3 | 81.5 |
| HC044 | G | 44 | 0.72 | 7.9 | 94 | 72 | 165 | 80 | 78.7 |
| HC046 | E | 46 | 0.89 | 6.3 | 85 | 93 | 169 | 82 | 81.3 |
| HC066 | G | 65 | 1 | 5.8 | 92 | 216 | 155 | 82 | 81.8 |
| HC069 | T | 68 | 0.84 | 7.3 | 94 | 95 | 187 | 82.2 | 82.3 |

| | | | | | | | | | |
|---|---|---|---|---|---|---|---|---|---|
| HC071 | S | 70 | 0.84 | 6 | 82 | 130 | 197 | 82.2 | 81.8 |
| HC075 | S | 74 | 0.9 | 7 | 93 | 126 | 177 | 82.2 | 82.2 |
| HC076 | K | 75 | 0.88 | 10.2 | 93 | 129 | 224 | 82 | 82.2 |
| HC077 | N | 76 | 0.74 | 5.4 | 94 | 64 | 139 | 81.2 | 80.9 |
| HC084 | N | 82A | 0.79 | 6.7 | 95 | 110 | 151 | 81.9 | 82 |
| HC085 | S | 82B | 0.77 | 6.9 | 94 | 86 | 134 | 82.3 | 82.3 |
| HC087 | R | 83 | 0.71 | 7 | 93 | 96 | 193 | 82.2 | 82 |
| HC112 | Q | 105 | 0.85 | 8.8 | 96 | 76 | 168 | 82 | 82.1 |
| HC119 | S | 112 | 0.72 | 4.9 | 93 | 91 | 136 | 79.5 | 79.2 |
| HC120 | S | 113 | 0.7 | 6 | 78 | 105 | 137 | 82.4 | 82.3 |
| HC122 | S | 115 | 1 | 6.2 | 92 | 90 | 155 | 82 | 82.1 |
| HC124 | K | 117 | 0.82 | 4.4 | 92 | 108 | 217 | 81.8 | 81.6 |
| HC127 | S | 120 | 0.74 | 7.2 | 88 | 78 | 214 | 82.3 | 81.9 |
| HC137 | S | 130 | 0.97 | 7.2 | 88 | 72 | 188 | 82.2 | 81.2 |
| HC139 | S | 134 | 1 | 7.2 | 93 | 84 | 163 | 82.3 | 82 |
| HC141 | G | 136 | 0.92 | 5 | 94 | 114 | 177 | 82.3 | 82.3 |
| HC142 | T | 137 | 0.92 | 8 | 95 | 91 | 148 | 82.4 | 81.4 |
| HC151 | D | 146 | 0.76 | 3 | 49 | 94 | 138 | 68.8 | 69.9 |
| HC158 | T | 153 | 0.71 | 5.5 | 95 | 97 | 136 | 82.2 | 81.7 |
| HC163 | S | 163 | 0.99 | 17.7 | 94 | 74 | 177 | 82 | 82.4 |
| HC164 | G | 164 | 0.99 | 4.4 | 94 | 106 | 144 | 80.9 | 80.3 |
| HC168 | S | 168 | 1 | 4.6 | 93 | 125 | 94 | 82.2 | 82.2 |
| HC179 | S | 180 | 0.98 | 2.8 | 91 | 117 | 125 | 82 | 81.8 |
| HC180 | S | 182 | 0.73 | 2.8 | 92 | 161 | 158 | 80.9 | 80.3 |
| HC181 | G | 183 | 0.97 | 3.7 | 97 | 126 | 160 | 81.5 | 82.4 |
| HC194 | S | 196 | 0.98 | 4.6 | 90 | 107 | 90 | 82.2 | 82 |
| HC197 | G | 199 | 0.77 | 4.2 | 92 | 166 | 191 | 82.5 | 82.4 |
| HC198 | T | 200 | 0.94 | 4 | 94 | 122 | 134 | 82.2 | 81.9 |
| HC202 | I | 207 | 0.73 | 5 | 35 | 72 | 157 | 81.9 | 81.2 |
| HC206 | N | 211 | 0.92 | 5.7 | 94 | 110 | 144 | 82.5 | 81.8 |
| HC211 | N | 216 | 1 | 10.1 | 85 | 74 | 192 | 82.2 | 82.3 |
| HC213 | K | 218 | 0.99 | 6.7 | 91 | 89 | 126 | 82.3 | 82.1 |
| HC222 | S | 229 | 0.88 | 7.8 | 91 | 113 | 128 | 82.5 | 82.5 |

**Table 2: Labeling, stability, and affinity of 23 less accessible methionine substitutions in model αGFP-Fab in trastuzumab scaffold.**

| Site | Residue | Kabat | ASA | Yield (mg/L) | $K_d$ (pM) | Thermo-stability°C | % Labeling at 5x | % Labeling at 20x | % Stability at 37°C |
|---|---|---|---|---|---|---|---|---|---|
| LC022 | T | 22 | 0.36 | 11.9 | 770 | 82.3 | 92 | - | 42 |
| LC063 | S | 63 | 0.52 | 10.8 | 190 | 81.9 | 76 | - | 59 |
| **LC066** | **R** | **66** | **0.40** | **6.9** | **780** | **82.7** | **68** | **91** | **99** |
| LC069 | T | 69 | 0.83 | 21.8 | 450 | 82.7 | 55 | 84 | 62 |
| LC070 | D | 70 | 0.40 | 0.4 | 230 | 82.8 | - | - | - |
| LC072 | T | 72 | 0.56 | 0.5 | 270 | 82.2 | - | - | - |
| **LC074** | **T** | **74** | **0.44** | **31.2** | **820** | **81.4** | **47** | **83** | **96** |
| LC147 | Q | 147 | 0.48 | 25.5 | 490 | 82.5 | 61 | 85 | 49 |
| LC148 | W | 148 | 0.10 | 2.0 | 630 | 75.6 | 0 | - | - |
| LC149 | K | 149 | 0.27 | 3.2 | 520 | 82.3 | 55 | - | 88 |
| LC192 | Y | 192 | 0.12 | 0.5 | 290 | 78.2 | - | - | - |
| LC193 | A | 193 | 0.46 | 13.3 | 430 | 81.8 | 0 | - | - |
| LC195 | E | 195 | 0.49 | 49.7 | 430 | 82.4 | 70 | 87 | 55 |
| LC208 | S | 208 | 0.57 | 5.9 | 380 | 82.4 | 0 | - | - |
| HC018 | L | 18 | 0.46 | 33.2 | 650 | 82.6 | 0 | - | - |
| HC021 | S | 21 | 0.54 | 36.7 | 530 | 82.3 | 52 | 78 | 87 |
| HC025 | S | 25 | 0.42 | 17.2 | 800 | 82.7 | 81 | - | 65 |
| HC082 | Q | 81 | 0.67 | 0.0 | - | - | - | - | - |
| HC201 | Y | 206 | 0.13 | 22.4 | 720 | 81.0 | 0 | - | - |
| HC204 | N | 209 | 0.47 | 5.1 | 730 | 81.7 | 84 | - | 46 |
| HC212 | T | 217 | 0.26 | 23.7 | 18200 | 81.8 | 62 | 83 | 95 |
| HC215 | D | 220 | 0.49 | 21.3 | 550 | 80.8 | 85 | - | 55 |
| HC217 | K | 222 | 0.64 | 20.5 | 530 | 82.3 | 62 | 84 | 60 |

**Supplementary Table 3: Alignment, labeling, and stability of chosen Fc sites.**

| Site | Residue | Kabat | Aligned Fab site | % labeling | % stability |
|---|---|---|---|---|---|
| HC262 | V | 275 | HC.S21 | 86 | 100 |
| HC292 | R | 309 | LC.R66 | 80 | 74 |
| HC307 | T | 326 | LC.T74 | 0 | - |
| HC382 | E | 407 | LC.K149 | 61 | 90 |
| HC437 | T | 468 | HC.T212 | 0 | - |

**Supplementary Methods:**

*Flow cytometry*

Cells were lifted with Versene (0.04% EDTA, PBS pH 7.4 Mg/Ca free), and subsequently blocked with flow cytometry buffer (PBS, pH 7.4, 3% BSA). Fabs or IgGs were added to cells for 30 min on ice. Antibodies were detected with addition of Alexa Fluor 647 AffiniPure F(ab')$_2$ conjugate (Jackson Immuno Researech, 1:50). Cells were extensively washed and fluorescence in the APC channel was quantified using a FACSCanto II (BD Biosciences). All flow cytometry data analysis was performed using FlowJo software.

*Size Exclusion Chromatography*

SEC analysis was performed using an Agilent HPLC 1260 Infinity II LC System using an AdvanceBio SEC column (300Å, 2.7μm, Agilent). Each analyte was injected at 5-10μM and run with a constant mobile phase of 0.15 M sodium phosphate for 15 minutes. Fluorescence (excitation 285nm, emission 340nm) and absorbance was measured.

**Supplementary Note:**

*Synthesis of azide-piperidine oxaziridine*

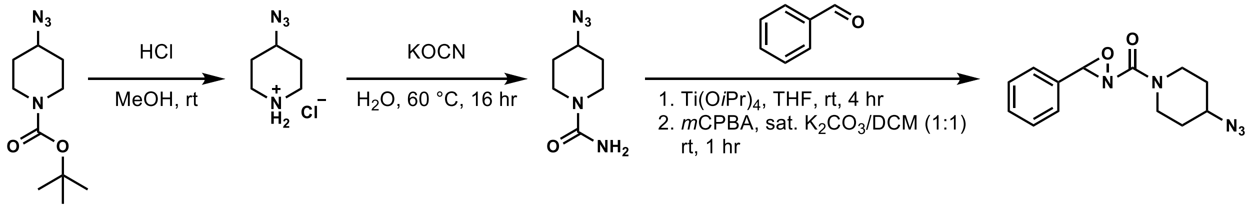


tert-butyl 4-azidopiperidine-1-carboxylate was synthesized using a previous published procedure[1].

tert-butyl 4-azidopiperidine-1-carboxylate (10 mmol) was dissolved in MeOH (20 mL) and 4M HCl in 1,4-dioxane (22 mmol, 2.2 equiv.) was added dropwise and the solution was stirred overnight at room temperature. The solution was concentrated *in vacuo* and a colorless solid formed after washed with diethyl ether (quantitative yield). The crude material was taken on without further purification.

4-azidopiperidinium chloride (10 mmol) was dissolved in water (10 mL) and KOCN (30 mmol, 3 equiv.) was added in one portion. The reaction was stirred at 60 °C for 12 hours and a colorless precipitant formed. The precipitant (4-azidopiperidine-1-carboxamide) was filter, washed with diethyl ether, and was used in the next reaction without any further purification (quantitative yield).

THF (30 mL) was added to 4-azidopiperidine-1-carboxamide (1.7 g, 10 mmol, 1 equiv.) and benzaldehyde (1.2 mL, 12 mmol, 1.2 equiv.) was subsequently added. Ti(O*i*Pr)$_4$ (4.1 mL, 14 mmol, 1.4 equiv.) was added and the reaction stirred at room temperature for 4 hours. The solution was concentrated under reduced pressure and was used immediately.

mCPBA (75%, 6.9 g, 30 mmol, 3 equiv.) was added to a 1:1 mixture of DCM:sat. $K_2CO_3$ (0.125 M) and was stirred for 10 minutes. Condensation residue (1 equiv.) was dissolved in DCM (1 M) and was added slowly to the mCPBA mixture. The mixture was vigorously stirred for 1 hour and was diluted with $H_2O$ (3x the volume). The biphasic solution was extracted with DCM (3x) and the organic layers combined, washed with brine (1x), dried over $Na_2SO_4$, and was concentrated under reduced pressure. The concentrate was purified using column chromatography (1% $Et_2O$ in DCM) to afford the oxaziridine as a colorless oil (891 mg, 33% yield).

**$^1$H NMR:** (500 MHz, $CDCl_3$) δ 7.53 – 7.38 (m, 5H), 5.23 (s, 1H), 4.17 (dt, *J* = 14.0, 5.1 Hz, 1H), 4.04 – 3.90 (m, 1H), 3.78 (dtd, *J* = 24.2, 7.4, 3.7 Hz, 1H), 3.68 (ddd, *J* = 13.4, 8.3, 3.8 Hz, 1H), 3.46 (dddd, *J* = 22.9, 13.1, 8.5, 3.6 Hz, 1H), 3.27 (ddd, *J* = 13.2, 9.2, 3.5 Hz, 1H), 2.02 – 1.86 (m, 2H), 1.69 (dddd, *J* = 35.1, 17.7, 8.8, 4.2 Hz, 2H).

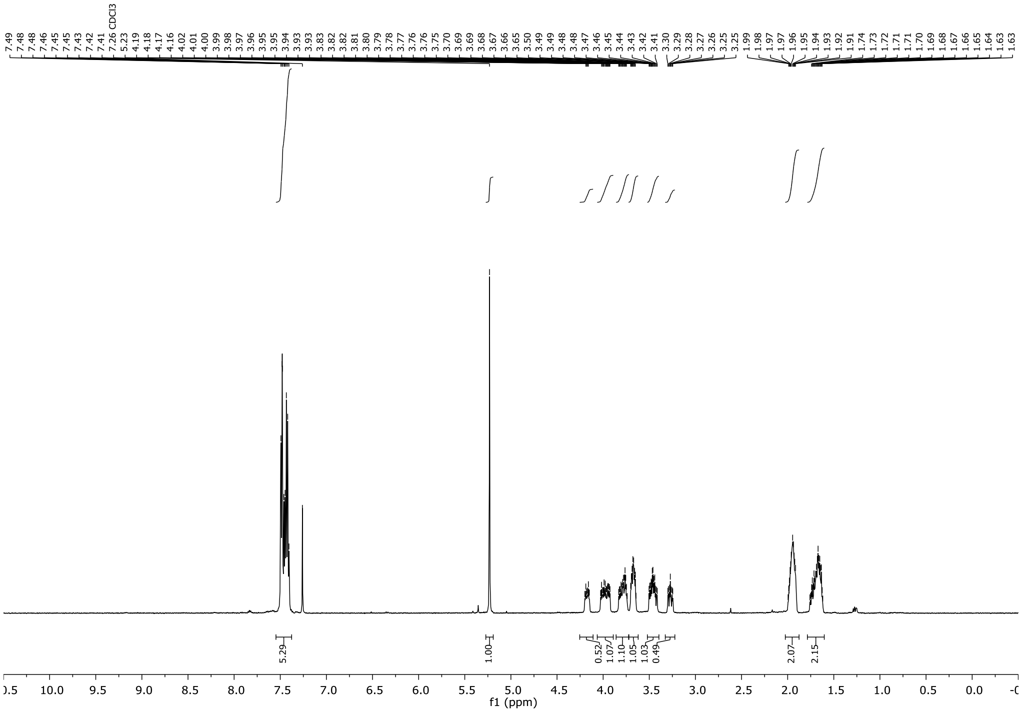


**$^{13}$C NMR:** (126 MHz, $CDCl_3$) δ 160.73, 160.71, 132.86, 132.83, 130.94, 128.79, 128.02, 78.18, 78.15, 57.08, 56.77, 42.65, 42.19, 41.60, 41.10, 30.98, 30.84, 30.41, 30.19.

**IR:** ($cm^{-1}$) 2931, 2089, 1686, 1429, 1248

**HRMS** (ESI) *m/z* calcd for $C_{13}H_{16}O_2N5$ $[M+H]^+$: 274.1299, found: 274.1296

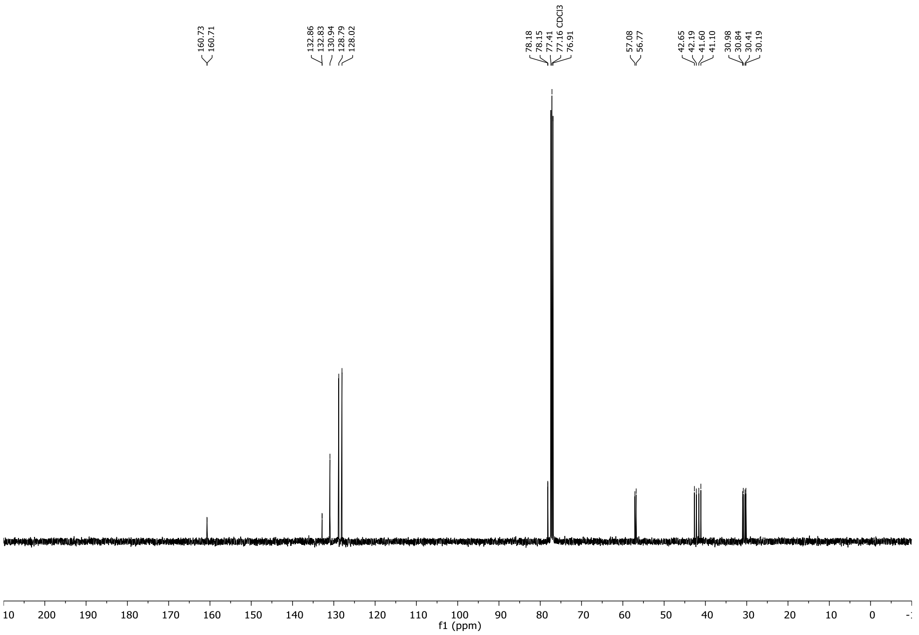

*Synthesis of adamantyl oxaziridine*

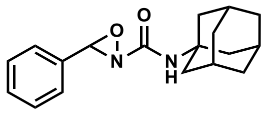

Adamantyl oxaziridine was synthesized using previously published procedures in 1% yield as a colorless solid[2].

**$^1$H NMR:** (600 MHz, $CDCl_3$) δ 7.48 – 7.43 (m, 3H), 7.41 (dd, $J$ = 8.0, 6.5 Hz, 2H), 5.82 (s, 1H), 4.96 (s, 1H), 2.14 – 2.09 (m, 3H), 2.01 (d, $J$ = 2.9 Hz, 6H), 1.69 (t, $J$ = 3.1 Hz, 6H)

**$^{13}$C NMR:** (151 MHz, $CDCl_3$) δ 160.31, 132.86, 130.99, 128.72, 128.09, 79.30, 52.16, 41.46, 36.29, 29.52

**IR:** ($cm^{-1}$) 3384, 2903, 2850, 2154, 1695

**HRMS** (ESI) *m/z* calcd for $C_{18}H_{22}O_2N_2$ $[M+H]^+$: 299.1754, found: 299.1752

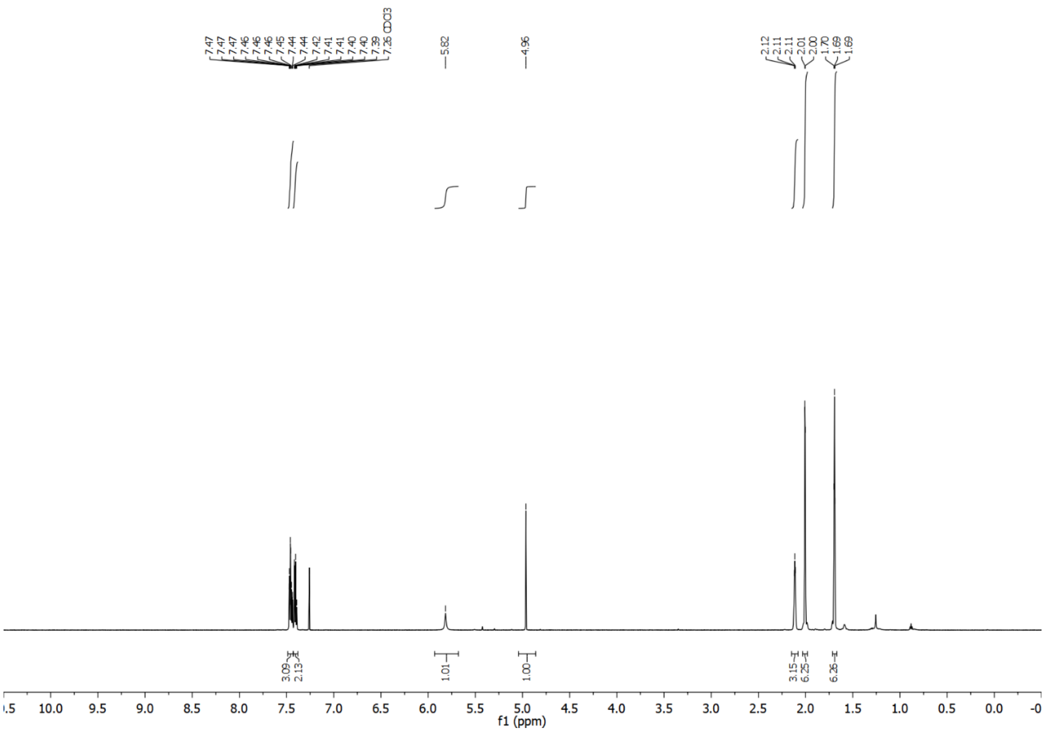

**References:**


1. Weisner, J. *et al.* Covalent-Allosteric Kinase Inhibitors. *Angew. Chem. Int. Ed. Engl.* **54**, 10313–6 (2015).
2. Christian, A. H. *et al.* A Physical Organic Approach to Tuning Reagents for Selective and Stable Methionine Bioconjugation. *J. Am. Chem. Soc.* jacs.9b04744 (2019). doi:10.1021/jacs.9b04744
